# IL-23 Licenses Pathogenic Th17 Transdifferentiation via STAT4-dependent Upregulation of IL-12Rβ2

**DOI:** 10.64898/2026.07.23.740320

**Authors:** Awalpreet S. Chadha, Asif Elahi, Blake F. Frey, Yoshiko Nagaoka-Kamata, Carlene L. Zindl, Trenton R. Schoeb, Caleb R. Glassman, K. Christopher Garcia, Casey T. Weaver

## Abstract

Th17 cells exhibit substantial developmental plasticity, transdifferentiating into pathogenic, interferon-γ (IFN-γ)–producing Th1-like effectors during chronic autoimmune inflammation. While both IL-23 and IL-12 are implicated in this process, how these structurally related cytokines coordinate to drive transdifferentiation remains unresolved. Here, we show that IL-23 acts as a developmental licensor rather than a terminal trigger, priming Th17 cells for transdifferentiation upon subsequent IL-12 encounter. IL-23 signals through STAT4 to upregulate the IL-12–specific receptor subunit IL-12Rβ2, lowering the signalling threshold for IL-12. This licensing can be conferred at both early and late stages of Th17 development and produces a temporal logic in which IL-23-mediated STAT4 signaling functionally arms Th17 cells to terminally transdifferentiate in response to IL-12. Using *Stat4* conditional knockouts and a structure-based IL-23 mutein that selectively abolishes IL-23–driven STAT4 signaling while preserving STAT3-dependent lineage maintenance, we genetically establish the IL-23–STAT4–IL-12Rβ2 axis as a discrete licensing module. Furthermore, using the adoptive Th17 transfer colitis model, we show that host-derived IL-23 and IL-12 are required for Th17-driven colitis, and the inflamed colonic environment reconstitutes transdifferentiation capacity in classically “non-pathogenic” Th17 cells. These findings reframe the pathogenic dichotomy of Th17 cells as one of cumulative cytokine exposure rather than fixed cellular fate, and reveal a stepwise architecture of Th17 plasticity in which IL-23 licensing and IL-12 triggering operate as functionally distinct events.

**Significance Statement:** Th17 cells protect mucosal barriers but can convert into pathogenic, IFN-γ-producing effectors that drive inflammatory bowel disease (IBD). Although IL-23 is essential for this conversion, the underlying molecular mechanism remains elusive. We show that IL-23 does not directly trigger transdifferentiation but instead licenses Th17 cells for this conversion, during both early and late stages of their development. By signaling through STAT4, IL-23 upregulates the IL-12–specific receptor subunit IL-12Rβ2 on Th17 cells, rendering them responsive to subsequent IL-12 exposure. Our findings reframe the classical “pathogenic” versus “non-pathogenic” Th17 distinction and provide a mechanistic basis for the clinical efficacy of IL-23–targeted therapies in IBD.

## Introduction

CD4⁺ T-helper 17 (Th17) cells are essential for host defense against extracellular pathogens at mucosal barriers but also possess substantial developmental plasticity that can drive severe autoimmune pathology, including inflammatory bowel disease (IBD)(1). During chronic mucosal inflammation, committed Th17 cells frequently extinguish IL-17A production and transdifferentiate into pathogenic, IFN-γ–producing Th1-like effectors — a process indispensable for disease in adoptive Th17 transfer models of colitis and central to encephalitogenic disease in experimental autoimmune encephalomyelitis (EAE)(2, 3). The cytokines that govern this transdifferentiation belong to the IL-12 family. IL-23 (p19/p40) and IL-12 (p35/p40) share a common p40 subunit and signal through heterodimeric receptor complexes that share IL-12Rβ1 but pair with distinct cytokine-specific chains — IL-23R for IL-23 and IL-12Rβ2 for IL-12 — that drive STAT3-dominant and STAT4-dominant signaling, respectively (4, 5). How these structurally related cytokines coordinate to drive Th17 transdifferentiation has remained mechanistically unclear.

IL-23 has historically been characterized as a survival and stabilization factor that sustains the Th17 lineage through STAT3-driven survival programs (4, 6–8). More recent single-cell transcriptomic studies have refined this view, identifying IL-23R signaling as a critical determinant of Th17 transcriptional state and a driver of the homeostatic-to-effector transition (9, 10). Yet how IL-23 signaling produces the specific molecular shift toward IFN-γ production has not been defined. IL-12, conversely, is indispensable for imprinting the Th1 program through STAT4, but its physiological availability is tightly constrained, particularly in mucosal tissues (11, 12). The combination of IL-23’s incompletely understood role in driving phenotypic Th17 conversion and IL-12’s restricted bioavailability raises the possibility that Th17 transdifferentiation requires a coordinated mechanism in which one cytokine prepares the cell to respond to the other.

Here, we define the molecular logic of this coordination. Using an antigen-specific SMARTA TCR-transgenic system, we demonstrate that IL-23 functions as a developmental licensor during both early and late Th17 development, actively licensing cells for pathogenic transdifferentiation without serving as the definitive trigger. We show that IL-23 stabilizes the Th17 phenotype while simultaneously signaling through STAT4 to upregulate IL-12Rβ2, thereby lowering the cellular activation threshold for IL-12 and enabling subsequent IL-12-driven induction of IFN-γ and T-bet. We genetically dissect the underlying IL-23–STAT4–IL-12Rβ2 axis using *Stat4* conditional knockouts and a structure-based IL-23 mutein that selectively abolishes IL-23–driven STAT4 signaling while preserving STAT3-dependent lineage maintenance, allowing us to isolate the licensing function of IL-23 from its lineage-stabilizing role. Finally, using *in vivo* competitive co-transfer models of Th17 transfer colitis, we show that both IL-12Rβ2 expression on Th17 cells and host-derived IL-23 and IL-12 are essential for transdifferentiation and disease induction, and that the inflamed colonic environment reconstitutes transdifferentiation capacity in classically polarized “non-pathogenic” Th17 cells. Together, these findings establish a stepwise model of Th17 plasticity in which IL-23 functionally arms Th17 cells to execute a pathogenic effector program upon subsequent IL-12 encounter.

## Results

### IL-23 licenses Th17 cells for IL-12–driven transdifferentiation at early and late stages of development

To define the developmental requirements for Th17-Th1 transdifferentiation, we used an antigen-specific SMARTA TCR-transgenic system to avoid confounders associated with polyclonal, nonspecific *in vitro* expansion. Naïve CD4⁺ T cells from IL-17F reporter (*Il17f^Thy1.1^*) mice crossed to SMARTA (*Il17f^Thy1.1^*·SMARTA) mice were polarized under classical Th17 conditions (TGF-β plus IL-6; “T+6”) or pathogenic Th17 conditions (IL-1β, IL-6, and IL-23; “1+6+23”), and sort-purified on the basis of Thy1.1 (IL-17F) expression to exclude non-Th17 cells (2, 13, 14). Sorted cells were restimulated with cognate gp61 peptide and splenic feeders in the presence of added T+6, IL-23, or titrated IL-12 (**Fig. 1A**). Consistent with prior work, T+6 and IL-23 each maintained a Th17 phenotype, whereas IL-12 drove rapid Th1-like transdifferentiation (**Fig. 1B**) (8, 15). Notably, prior IL-23 exposure markedly lowered the IL-12 threshold for this transition: 1+6+23–primed cells generated robust IFN-γ responses at IL-12 concentrations 100–1,000-fold lower than those required to elicit comparable conversion of T+6 cells (**Fig. 1B and Fig. S1A and B**). Thus, IL-23 exposure during early Th17 development heightens their sensitivity to IL-12–driven transdifferentiation.

**Figure 1.**
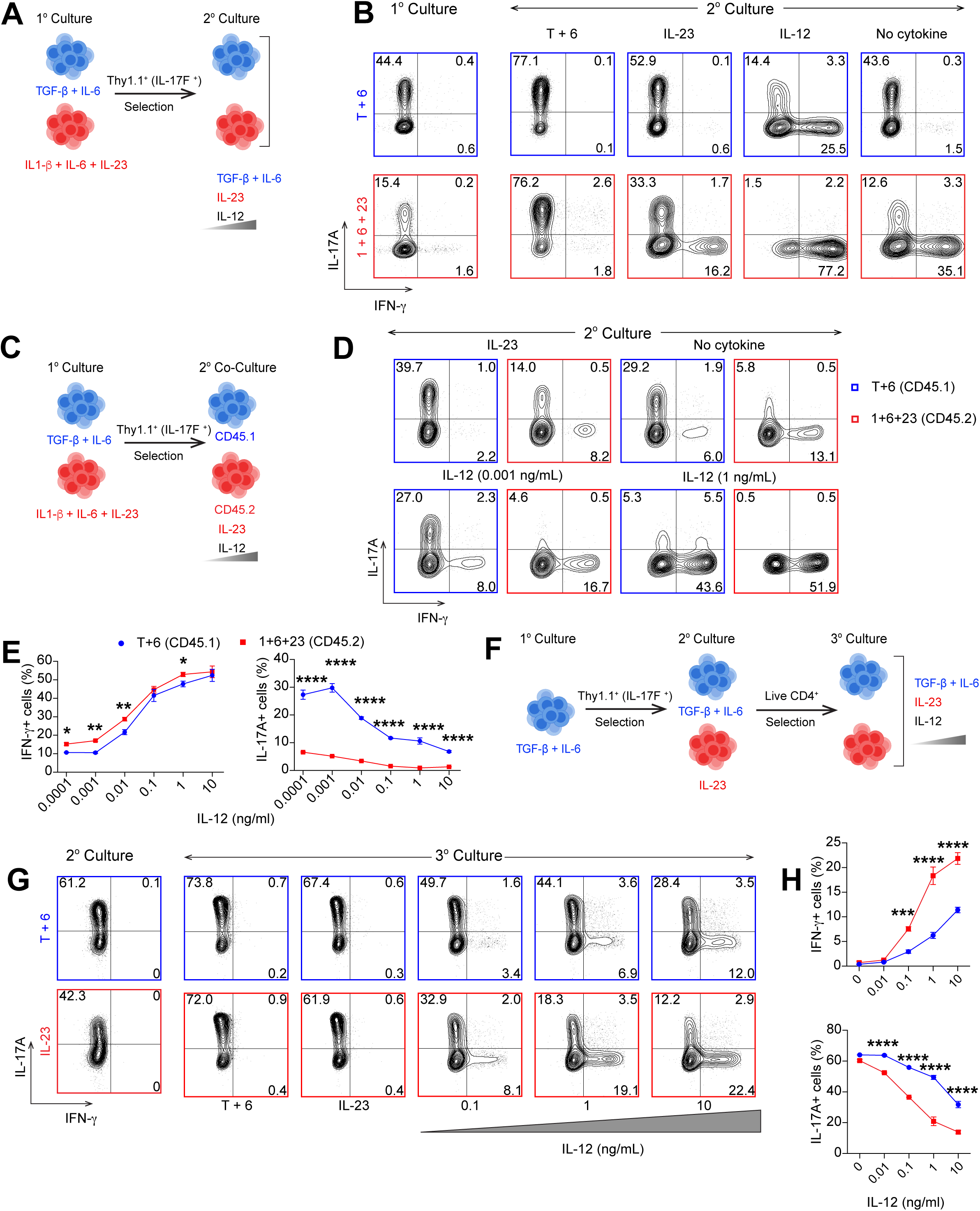
IL-23 licenses Th17 cells for IL-12–driven transdifferentiation at early and late stages of development. (*A*) Schematic of primary Th17 polarization and secondary culture. Naïve CD4⁺ T cells from *Il17f^Thy1.1^* ·SMARTA mice were cultured with irradiated splenic feeders under classical Th17 (2.5 ng/mL TGF-β + 10 ng/mL IL-6; T+6) or pathogenic Th17 (20 ng/mL IL-1β + 10 ng/mL IL-6 + 10 ng/mL IL-23; 1+6+23) polarizing conditions for 5 days with 1.25 μg/mL gp61 peptide. IL-17F^Thy1.1+^ cells were sort-purified and restimulated with gp61 peptide and irradiated feeders in T+6, IL-23, titrated IL-12, or no cytokine for an additional 5 days. (*B*) Representative FACS plots of IL-17A and IFN-γ expression in T+6– and 1+6+23–primed cells after secondary culture in the indicated conditions. Plots gated on live CD4⁺ T cells. (*C*) Schematic of competitive co-culture: IL-17F^Thy1.1+^-sorted T+6 (CD45.1) and 1+6+23 (CD45.2) cells were co-cultured 1:1 across a graded IL-12 dose range. (*D*) Representative FACS plots of co-cultured T+6 and 1+6+23 cells at the indicated IL-12 doses. (*E*) Frequencies of IFN-γ⁺ and IL-17A⁺ cells in each co-cultured population across IL-12 doses (mean ± SD). (*F*) Schematic of late-licensing tertiary culture: T+6–primed IL-17F^Thy1.1+^ cells were maintained in T+6 or IL-23 during secondary culture, sorted on live CD4⁺ cells, then challenged with titrated IL-12 in tertiary culture. (*G*) Representative FACS plots of IL-17A and IFN-γ expression after secondary culture and following tertiary IL-12 challenge across the indicated dose range. (*H*) Frequencies of IFN-γ⁺ and IL-17A⁺ cells across IL-12 doses in tertiary culture (mean ± SD). Data are representative of three independent experiments with three biological replicates per condition. *P < 0.05, **P < 0.01, ***P < 0.001, ****P < 0.0001.

To determine if this heightened sensitivity was a stable, cell-intrinsic trait or a consequence of paracrine signaling, we performed competitive co-cultures of congenically marked T+6 (CD45.1) and 1+6+23 (CD45.2) Thy1.1^+^ Th17 cells across a graded IL-12 dose range (**Fig. 1C**). Within the same microenvironment, each population retained its distinct IL-12 sensitivity profile: 1+6+23 cells efficiently acquired IFN-γ and extinguished IL-17A at low IL-12 doses, whereas co-cultured T+6 cells remained comparatively refractory (**Fig. 1D** and **E**), indicating that IL-23 establishes a cell-intrinsic licensing state that confers heightened IL-12 responsiveness on Th17 cells.

Th17 cells retain substantial developmental plasticity after lineage commitment and their fate is shaped by cytokines they subsequently encounter (8). We therefore asked whether IL-23 licensing is restricted to early Th17 differentiation or can also be conferred on cells already committed to Th17. *Il17f^Thy1.1^*T+6–primed Th17 cells were maintained in either T+6 or IL-23 during secondary culture and then challenged with titrated IL-12 in tertiary culture (**Fig. 1F**). Prolonged maintenance in IL-23 alone did not induce IFN-γ, instead serving as a lineage anchor that sustained robust IL-17A expression indistinguishable from T+6–maintained cells (**Fig. 1G**). Yet these two phenotypically similar populations responded very differently to IL-12: cells maintained in IL-23 cells underwent accelerated transdifferentiation, acquiring IFN-γ and extinguishing IL-17A compared to their T+6 counterparts (**Fig. 1G** and **H**). Collectively, these data suggest that IL-23 does not itself drive transdifferentiation but maintains a primed Th17 state competent to respond to IL-12, and this licensing can be acquired at both early and late stages of Th17 development.

### IL-23 licenses Th17 cells through STAT4-mediated upregulation of IL-12Rβ2

Building on our findings that sequential exposure of IL-23 followed by IL-12 accelerated Th17 transdifferentiation regardless of the developmental stage at which IL-23 was encountered, we next sought to define the receptor-level mechanism underlying this temporal logic. IL-12 and IL-23 receptor complexes share the signaling subunit IL-12Rβ1 but pair with distinct chains — IL-12Rβ2 for IL-12 and IL-23R for IL-23 — that drive cytokine-specific STAT4- and STAT3-dominant signaling, respectively (5, 16, 17). This shared architecture predicts that simultaneous exposure to IL-12 and IL-23 should generate competition at the level of receptor assembly. To test this, we examined the effect of simultaneous exposure of IL-23 and IL-12 on Th17 plasticity. Unlike sequential exposure, concurrent exposure to IL-12 and high-dose IL-23 (10 ng/mL) restricted transdifferentiation, resulting in cells of a Th17 state with higher IL-17A and lower IFN-γ production, whereas low-dose IL-23 (1 ng/mL) produced an intermediate phenotype relative to IL-12 alone (**Fig. 2A-C**). Thus, concurrent IL-23 signaling counterbalances IL-12–driven transdifferentiation in a dose-dependent manner.

**Figure 2.**
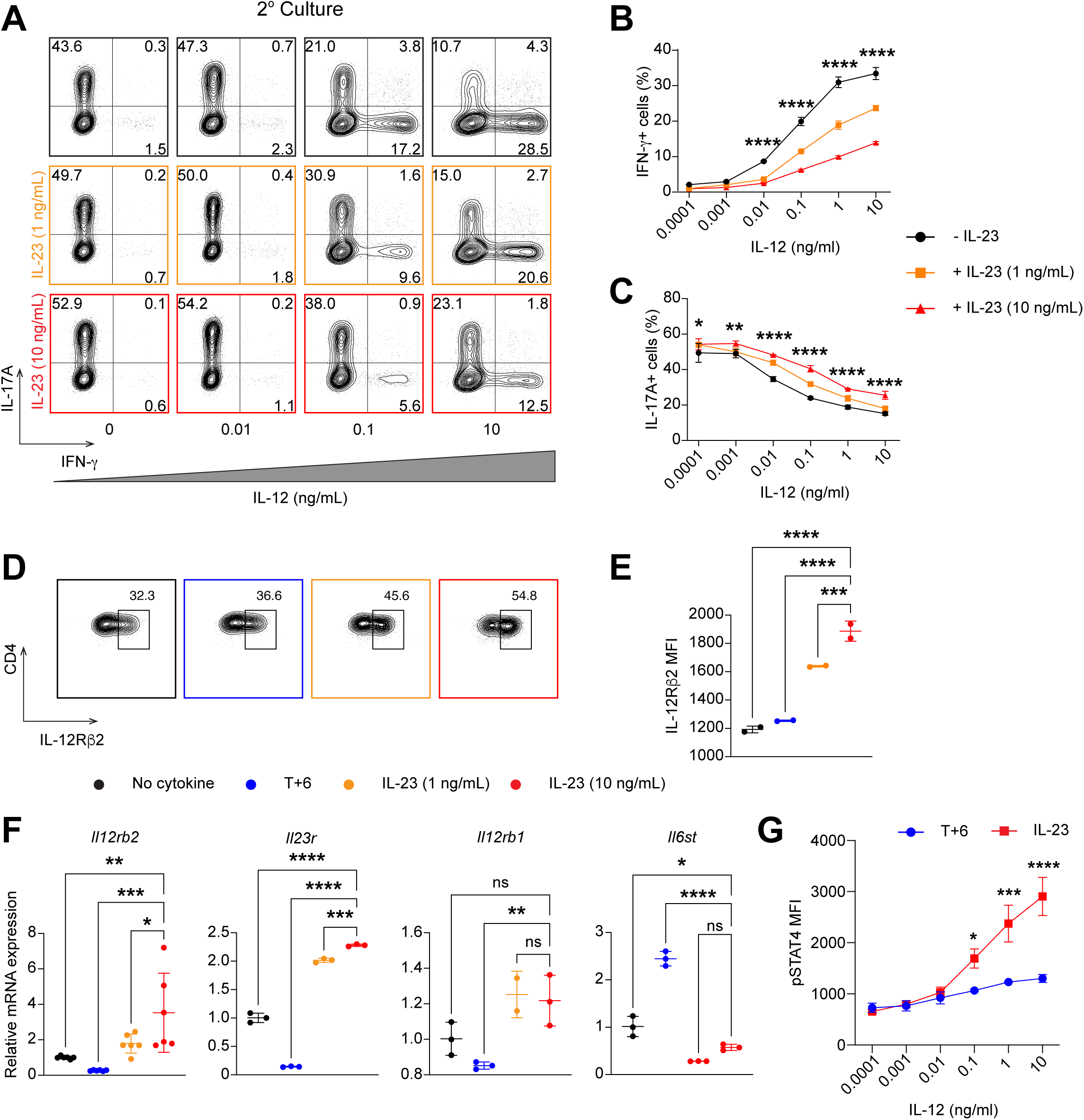
IL-23 licenses Th17 cells through STAT4-mediated upregulation of IL-12Rβ2. (*A*) Representative FACS plots of IL-17A and IFN-γ expression in T+6–primed IL-17F^Thy1.1+^ Th17 cells cultured with titrated IL-12 alone or concurrently with low (1 ng/mL) or high (10 ng/mL) IL-23 for 5 days. Plots gated on live CD4⁺ T cells. (*B* and *C*) Frequencies of IFN-γ⁺ (*B*) and IL-17A⁺ (*C*) cells across IL-12 doses with concurrent low- or high-dose IL-23 exposure (mean ± SD). (*D*) Representative FACS plots of surface IL-12Rβ2 expression on Th17 cells maintained in secondary culture with no cytokine, T+6 or low- and high-dose IL-23 for 5 days. (*E*) IL-12Rβ2 MFI across IL-23 doses (mean ± SD). (*F*) *Il12rb2, Il23r, Il12rb1* and *Il6st* transcript quantification by qRT-PCR in Th17 cells maintained with no cytokine, T+6 or low- and high-dose IL-23, normalized to *Gapdh* (mean ± SD). (*G*) pSTAT4 MFI in IL-23–maintained and T+6–maintained Th17 cells in secondary culture following 45-min acute re-challenge with titrated IL-12 (mean ± SD). Data are representative of two to three independent experiments with three biological replicates per condition. *P < 0.05, **P < 0.01, ***P < 0.001, ****P < 0.0001.

This antagonism raised a mechanistic question: if IL-23 and IL-12 oppose one another when present together, how does prior IL-23 exposure produce heightened IL-12 responsiveness? Although IL-23 predominantly signals through pSTAT3, it also generates low levels of pSTAT4, which can bind the *Il12rb2* locus and induce its transcription (4, 18). We therefore hypothesized that IL-23 priming resolves this conflict by upregulating the IL-12–specific subunit IL-12Rβ2 through STAT4 signaling, thereby enhancing sensitivity to IL-12 and shifting the balance in favor of IL-12 signaling upon subsequent IL-12 encounter. Flow cytometric and transcriptional analyses revealed that Th17 cells maintained in IL-23 exhibited dose-dependent increases in *Il12rb2* transcripts and surface IL-12Rβ2 protein relative to T+6 and TCR/co-stimulation only– maintained controls, with a concomitant increase in *Il23r* but no change in *Il12rb1* expression (**Fig. 2D-F**). Notably, since IL-12Rβ2 increased in a dose-dependent manner above the cytokine-free control, this reflects active induction by IL-23 rather than mere relief of TGF-β– mediated suppression (19). This selective upregulation of *Il12rb2* and *Il23r* was also observed when IL-23 was present during initial Th17 priming (**Fig. S1C**). Conversely, *Il6st* (encoding gp130) was elevated in T+6– relative to IL-23–maintained Th17 cells in secondary culture, with a similar pattern observed between T+6– and 1+6+23–primed cells during primary culture (**Fig. 2F** and **Fig. S1C**). The presence of TGF-β was therefore associated with reciprocal regulation of IL-12– and IL-6–family receptor subunits — reduced *Il12rb2* and increased *Il6st* — opposing the IL-23–driven upregulation of *Il12rb2*.

To determine whether this receptor reconfiguration translated into amplified intracellular signaling, we re-challenged IL-23–maintained and T+6–maintained Th17 cells in secondary culture with titrated IL-12 doses and measured STAT4 phosphorylation (**Fig. 2G**). IL-23– maintained cells exhibited markedly elevated pSTAT4 across stimulatory IL-12 doses, demonstrating that IL-12Rβ2 upregulation functionally amplifies IL-12 signal transduction.

Together, these findings suggest a molecular basis for IL-23 licensing. When present simultaneously with IL-12, IL-23 competes for shared receptor components and stabilizes a Th17 state, whereas prior IL-23 exposure circumvents this competition by selectively upregulating both cytokine-specific receptor subunits: *Il23r* upregulation reinforces the Th17 lineage anchor, while parallel *Il12rb2* upregulation shifts receptor stoichiometry to favor IL-12 responsiveness. The net effect is that a Th17 cell exposed to IL-23 maintains its lineage identity while becoming competent to respond to a subsequent IL-12 encounter — establishing a temporal logic in which preceding IL-23 exposure licenses Th17 cells for subsequent IL-12– driven transdifferentiation into ex-Th17 Th1 cells.

### STAT4 signaling and IL-12Rβ2 expression are required for Th17 licensing and pathogenic transdifferentiation

In view of our findings that IL-23 upregulates IL-12Rβ2 with resulting amplification in pSTAT4 signaling upon IL-12 exposure, we next wanted to interrogate the role of STAT4 in this licensing axis. We first utilized *Stat4^fl/fl^Cd4^Cre^*mice, in which STAT4 is deleted specifically in CD4⁺ T cells. *Stat4*-deficient CD4⁺ T cells underwent normal Th17 polarization and maintenance, producing IL-17A at levels indistinguishable from *Cre^−^* controls (**Fig. 3A**). Upon IL-12 re-challenge in secondary culture, however, *Stat4*-deficient T cells failed to transdifferentiate, exhibiting near-complete loss of IFN-γ acquisition, pSTAT4 induction, and T-bet upregulation across all IL-12 doses (**Fig. 3B-D**). STAT4 is therefore strictly required for IL-12–driven transdifferentiation of Th17 cells.

**Figure 3.**
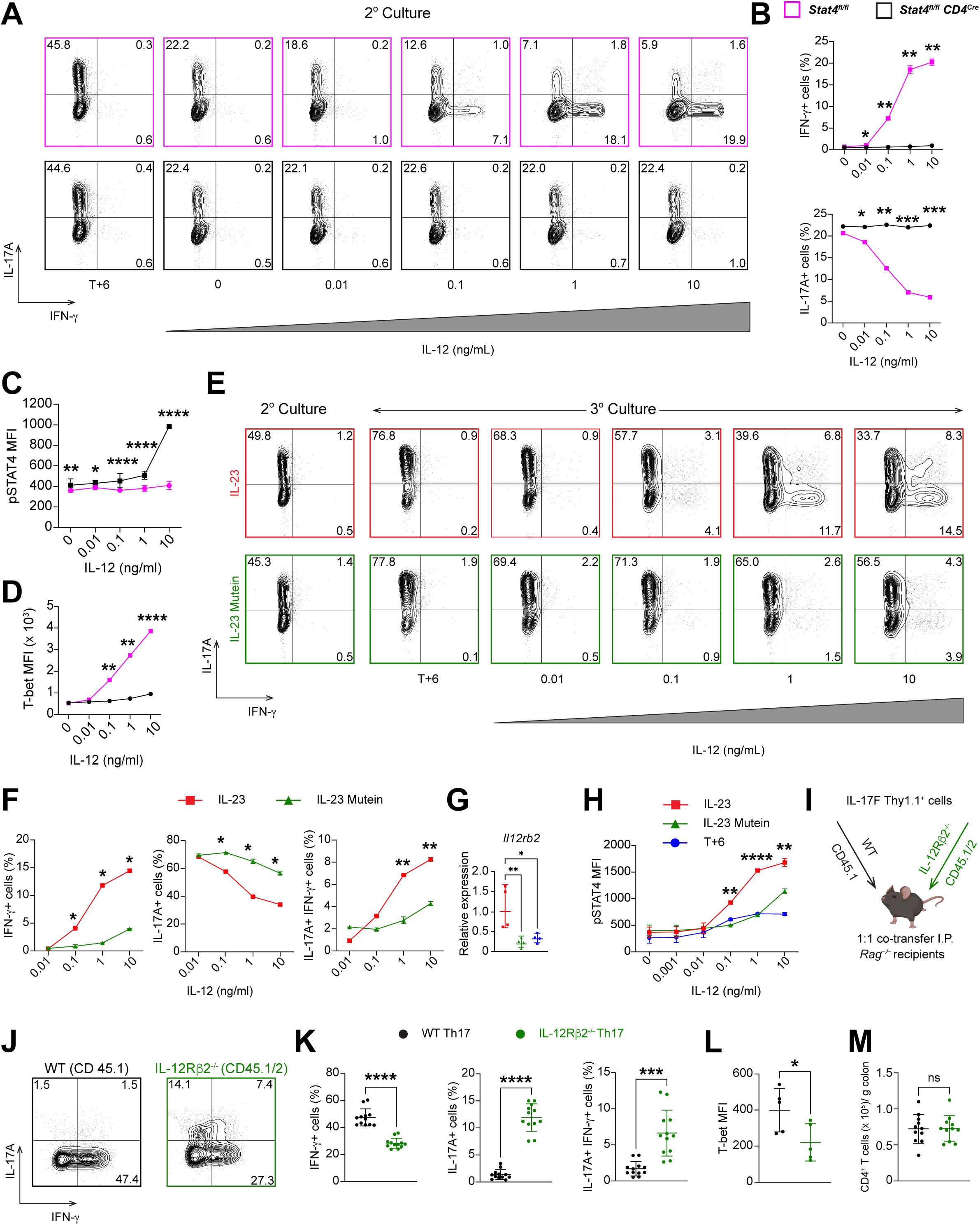
STAT4 signaling and IL-12Rβ2 expression are required for Th17 licensing and pathogenic transdifferentiation. (*A*) Representative FACS plots of IL-17A and IFN-γ expression in *Stat4^fl/fl^* (control) and *Stat4*^fl/fl^*Cd4*^Cre^ Th17 cells re-challenged with titrated IL-12 in secondary culture. Plots gated on live CD4⁺ T cells. (*B–D*) Frequencies of IFN-γ⁺ and IL-17A⁺ cells (*B*), pSTAT4 MFI (*C*), and T-bet MFI (*D*) across IL-12 doses in *Stat4*-sufficient and *Stat4*-deficient Th17 cells (mean ± SD). (*E*) Representative FACS plots of IL-17A and IFN-γ expression in IL-23– or IL-23 mutein–maintained Th17 cells after secondary culture and following tertiary challenge with T+6 or graded IL-12. (*F*) Frequencies of IFN-γ⁺, IL-17A⁺, and IL-17A⁺IFN-γ⁺ cells in IL-23– vs IL-23 mutein–maintained cells (mean ± SD). (*G*) *Il12rb2* transcript quantification in IL-23–, IL-23 mutein–, and T+6–maintained cells by qRT-PCR, normalized to *Gapdh* (mean ± SD). (*H*) pSTAT4 MFI in IL-23–, IL-23 mutein–, and T+6–maintained cells following 45-min acute IL-12 re-challenge (mean ± SD). (*I*) Schematic of competitive co-transfer: IL-17F^Thy1.1+^ -sorted T+6–polarized WT (CD45.1) and *Il17f^Thy1.1^*·*Il12rb2*^−/−^ (CD45.1/2) Th17 cells were co-transferred 1:1 into *Rag1^−/−^* recipients (4 x 10^5^ total cells per mouse). (*J*) Representative FACS plots of IL-17A and IFN-γ expression in WT and *Il12rb2*^−/−^ cells recovered from the CLP 4 weeks after transfer. (*K*) Frequencies of IFN-γ⁺, IL-17A⁺, and IL-17A⁺IFN-γ⁺ cells (mean ± SD). (*L*) T-bet MFI in recovered cells (mean ± SD). (*M*) Total CD4⁺ T cell recovery per gram of colonic tissue (mean ± SD). Data are representative of three (*in vitro*) or two (*in vivo*) independent experiments; in vivo data pooled from 10-12 mice per group. *P < 0.05, **P < 0.01, ***P < 0.001, ****P < 0.0001.

While these findings establish STAT4 as the central transducer of the transdifferentiation program, they do not distinguish whether STAT4 is required for IL-23–driven licensing (upstream IL-12Rβ2 upregulation) or only for IL-12-driven terminal signaling (downstream during transdifferentiation itself). In *Stat4*-deficient T cells, both functions are abolished simultaneously, leaving open the possibility that IL-23 upregulates IL-12Rβ2 through a STAT4-independent mechanism and that STAT4 is required only for IL-12 effector function. To selectively interrogate the licensing function of IL-23–driven STAT4 while preserving IL-12–mediated STAT4 signaling, we leveraged a structure-based IL-23 mutein engineered to dissect shared receptor usage (20). This mutein carries targeted substitutions in the p40 subunit (2xAla/3xAla/4xAla) that disrupt the p40–IL-12Rβ1 interface reducing IL-12Rβ1 recruitment, thereby attenuating STAT signaling relative to wild-type IL-23 while preserving IL-23R engagement through the unaltered p19 subunit (20). T+6 primed Th17 cells maintained in wild-type (WT) IL-23 or the IL-23 mutein during secondary culture were challenged with titrated IL-12 in tertiary culture. As anticipated, cells cultured with both WT IL-23 and IL-23 mutein maintained Th17 phenotype and pSTAT3 levels in secondary culture and produced equivalent amounts of IL-17 with T+6 during tertiary culture, confirming that the mutein supports STAT3-dependent lineage maintenance (**Fig. 3E**, **Fig. S2A**). Upon IL-12 exposure, however, WT IL-23–maintained cells underwent robust transdifferentiation, whereas mutein-maintained cells largely failed to acquire IFN-γ or extinguish IL-17A across the entire IL-12 dose range (**Fig. 3E** and **F**).

We next asked whether this functional defect reflected a failure of receptor licensing. Mutein-maintained cells failed to upregulate *Il12rb2*, displaying levels indistinguishable from T+6 controls (**Fig. 3G**). Consequently, upon acute IL-12 re-stimulation mutein-maintained cells exhibited severely blunted pSTAT4 compared to WT IL-23, mirroring unlicensed T+6 controls (**Fig. 3H**). Importantly, *Il12rb1* expression did not differ between WT IL-23 and IL-23 mutein maintained cells, while *Il23r* was elevated in both compared to T+6 controls, confirming preservation of STAT3 signaling by the mutein (**Fig. S2B**). Thus, the STAT4 signaling flux generated by IL-23, although weaker than its dominant STAT3 output, is sufficient to upregulate IL-12Rβ2 and license Th17 cells for subsequent IL-12 responsiveness.

Having genetically dissociated the licensing signal (IL-23–driven STAT4) from the lineage anchor (STAT3) and from the terminal trigger (IL-12–driven STAT4), we next tested whether IL-12Rβ2 is required to execute pathogenic transdifferentiation *in vivo*. To enable sorting of bona fide Th17 cells with or without IL-12Rβ2 expression, we crossed IL-17F reporter mice (*Il17f^Thy1.1^*) onto the *Il12rb2^−/−^*background (2). *Il12rb2*-deficient CD4^+^ T cells differentiated normally under T+6 conditions, producing equivalent IL-17A compared to wild-type (WT) controls (**Fig. S2C-D**), confirming that IL-12Rβ2 is dispensable for Th17 lineage commitment (2, 21). We then co-transferred congenically marked WT (CD45.1) and *Il12rb2^−/−^* (CD45.1/2) *Il17f^Thy1.1^* Th17 cells (both T+6–primed) at a 1:1 ratio into *Rag1^−/−^* recipients (**Fig. 3I**). Within an identical colonic inflammatory milieu, WT cells executed the pathogenic program, downregulating IL-17A and acquiring IFN-γ and T-bet, whereas *Il12rb2*-deficient Th17 cells were resistant to transdifferentiation, with markedly reduced IFN-γ⁺ and IL-17A⁺IFN-γ⁺ frequencies, lower T-bet expression, and sustained IL-17A production (**Fig. 3J-L**). Equivalent CD4⁺ T cell recovery from the colonic lamina propria (CLP) across genotypes excluded differential engraftment as a confounder (**Fig. 3M**).

Taken together, these findings establish a coherent molecular axis for IL-23 licensing: IL-23 activates STAT4 to upregulate IL-12Rβ2, IL-12Rβ2 expression is required to translate subsequent IL-12 encounter into amplified STAT4 signaling, and this amplified signal drives terminal Th1-like transdifferentiation. Genetic loss of any node — STAT4 itself, IL-23–driven STAT4 output, or IL-12Rβ2 expression on Th17 cells — selectively disrupts pathogenic transdifferentiation without compromising Th17 lineage maintenance, establishing the IL-23– STAT4–IL-12Rβ2 axis as a discrete, dissociable licensing module.

### Cumulative IL-23 and IL-12 exposure determines Th17 pathogenicity in vivo

Having established the molecular requirements for Th17 transdifferentiation, we next asked how prior IL-23 priming and ongoing host cytokine signaling might jointly determine pathogenic potential *in vivo*. Prior work in experimental autoimmune encephalomyelitis (EAE) established that 1+6+23–polarized Th17 cells transfer severe disease whereas T+6–polarized cells are largely non-encephalitogenic, a contrast that has long anchored the “pathogenic” versus “non-pathogenic” Th17 paradigm (13, 22, 23). To test whether this dichotomy holds in colitis, we adoptively transferred T+6 or 1+6+23 Thy1.1^+^ (IL-17F^+^) Th17 cells individually into *Rag1*^−/−^ recipients and tracked disease over five weeks (**Fig. 4A**). Contrary to the EAE precedent — and in agreement with our prior findings that T+6 cells potently induce colitis upon transfer — both populations induced equivalent wasting and colitis (**Fig. 4B and C**), with comparable frequencies of IFN-γ⁺, IL-17A⁺, and IL-17A⁺ IFN-γ⁺ cells, and equivalent T-bet expression among CD4⁺ T cells isolated from colonic lamina propria (CLP; **Fig. 4D-F**) (2, 8). There was also no difference in the myeloid populations isolated from CLP of the two groups (**Fig. S3A-C**). T+6 Th17 cells were, by these measures, indistinguishable from their IL-23–primed counterparts in the colitis setting. This equivalence suggested that the host colonic environment may itself supply the IL-23 signal necessary to license unprimed T+6 cells *in situ*, masking the cell-intrinsic differences observed *in vitro*.

**Figure 4.**
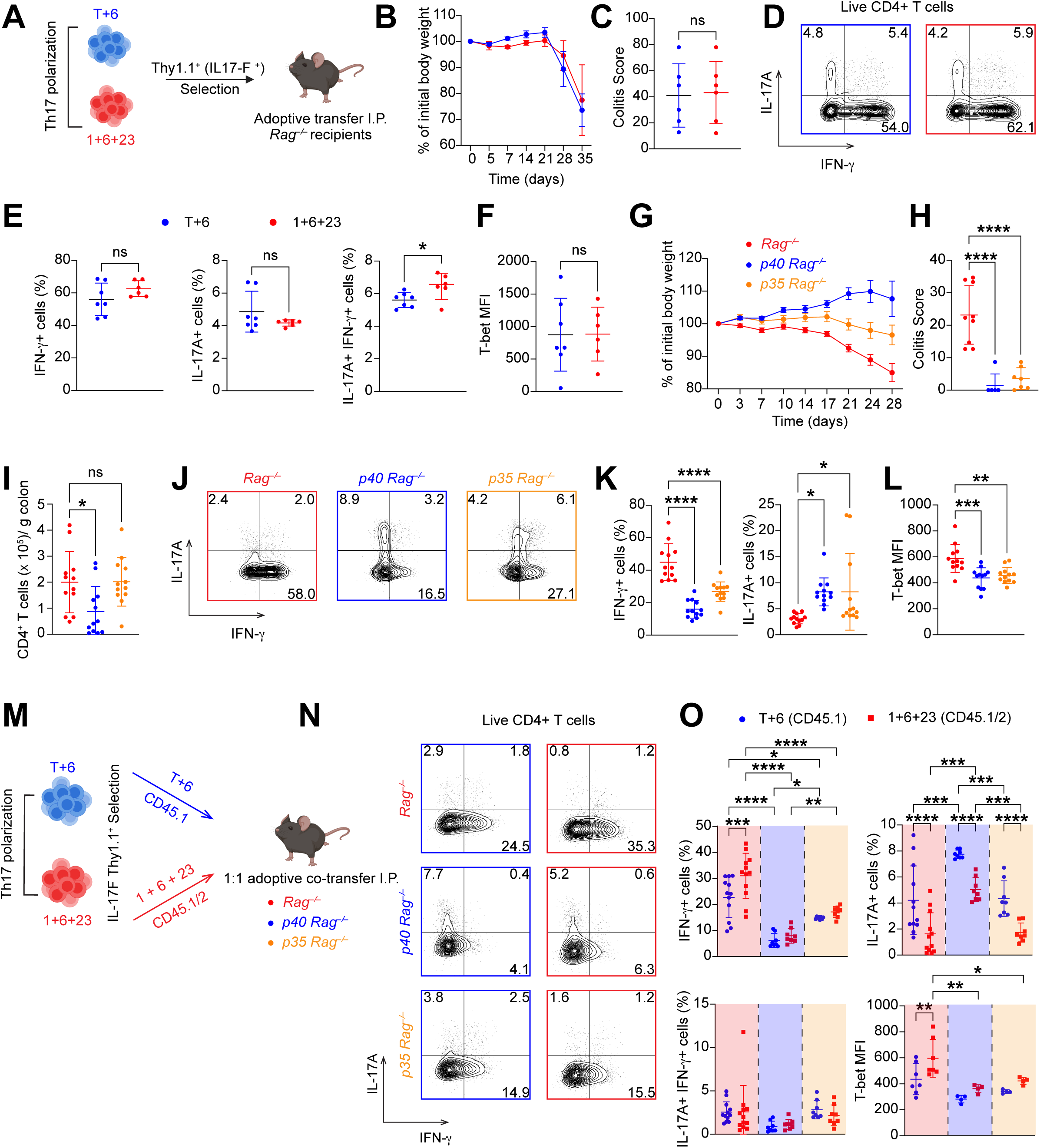
Cumulative host-derived IL-23 and IL-12 exposure determines pathogenic Th17 potential in vivo. (*A*) Schematic of adoptive transfer colitis: IL-17F^Thy1.1+^-sorted T+6– or 1+6+23–polarized Th17 cells were transferred separately into *Rag1^−/−^* recipients (4 x 10^5^ cells per mouse). (*B*) Weight loss expressed as a percentage of starting weight across 5 weeks post-transfer (mean ± SEM). (*C*) Histological scoring of colon sections at endpoint (mean ± SD). (*D*) Representative FACS plots of IL-17A and IFN-γ expression in T+6– and 1+6+23 transferred cells. (E) Frequencies of IFN-γ⁺ and IL-17A⁺ cells, IL-17A⁺IFN-γ⁺ cells, and T-bet MFI (*F*) in CD4⁺ T cells recovered from the CLP (mean ± SD). (*G*) Weight loss across 4 weeks post-transfer of T+6 Th17 cells into *Rag1*^−/−^, *Il12a*^−/−.^*Rag1*^−/−^ (p35 *Rag*^−/−^), or *Il12b*^−/−.^*Rag1*^−/−^ (p40 *Rag*^−/−^) recipients (mean ± SEM). (*H*) Histological scoring of colon sections at endpoint (mean ± SD). (*I*) Total CD4⁺ T cell recovery per gram of colonic tissue (mean ± SD). (*J*) Representative FACS plots of IL-17A and IFN-γ expression in CLP CD4⁺ T cells from each host. (*K* and *L*) Frequencies of IFN-γ⁺ and IL-17A⁺ cells (*K*) and T-bet MFI (*L*) in CLP CD4⁺ T cells from each host (mean ± SD). (*M*) Schematic of competitive co-transfer: T+6 (CD45.1) and 1+6+23 (CD45.1/2) Th17 cells were co-transferred 1:1 into *Rag1^−/−^*, p35 *Rag^−/−^*, or p40 *Rag^−/−^* recipients. (*N*) Representative FACS plots of IL-17A and IFN-γ expression in co-transferred T+6 and 1+6+23 cells across host backgrounds. (*O*) Frequencies of IFN-γ⁺, IL-17A⁺, and IL-17A⁺IFN-γ⁺ cells, and T-bet MFI in each co-transferred population across host backgrounds (mean ± SD). Only significant differences are shown. Data are pooled from two independent experiments with 6–12 mice per group. *P < 0.05, **P < 0.01, ***P < 0.001, ****P < 0.0001.

To test this directly, we transferred T+6–polarized Thy1.1^+^ (IL-17F^+^) Th17 cells into *Rag1*^−/−^ recipients, with or without deficiency of IL-12 (*Il12a*–deficient; *p35 Rag1*^−/−^), or both IL-12 and IL-23 (*Il12b*–deficient; *p40 Rag1*^−/−^). WT *Rag1*^−/−^ hosts developed rapid weight loss and severe colitis, whereas disease was markedly attenuated in both p35- and p40-deficient recipients (**Fig. 4G** and **H**). This attenuation was in part driven by differential T cell recovery: CD4⁺ T cell numbers in the CLP were reduced in p40-deficient recipients relative to p35-deficient and *Rag1*^−/−^ hosts (**Fig. 4I**), consistent with prior work establishing IL-23R signaling as a cell-intrinsic requirement for T cell accumulation and survival at the inflamed intestinal mucosa (7). In contrast, CD4⁺ T cell recovery in p35-deficient hosts was equivalent to *Rag1*^−/−^ recipients, indicating that the attenuated disease and reduced transdifferentiation observed in the absence of host IL-12 was not attributable to compromised T cell survival. Flow cytometric analysis of CLP CD4^+^ T cells confirmed that host cytokine availability dictated Th17 cell fate: Th17 cells robustly acquired IFN-γ, extinguished IL-17A, and upregulated T-bet in *Rag1*^−/−^ hosts but transdifferentiation was significantly reduced in both p35- and p40-deficient recipients (**Fig. 4J-L**). Notably, while both disease and transdifferentiation were markedly attenuated, neither was fully abolished in p35- or p40-deficient hosts, indicating that IL-12– and IL-23–independent pathways can partially sustain IFN-γ production and colonic pathology *in vivo*.

To directly compare the relative contributions of early versus late IL-23 exposure in Th17 transdifferentiation, and to dissociate IL-23 licensing from a subsequent IL-12 trigger, we co-transferred congenically marked T+6 (CD45.1) and 1+6+23 (CD45.1/2) Th17 cells at a 1:1 ratio into *Rag1*^−/−^, *p35 Rag1*^−/−^, or *p40 Rag1*^−/−^ hosts (**Fig. 4M**). In *Rag1^−/−^* recipients, 1+6+23 cells generated significantly more IFN-γ⁺ cells than co-transferred T+6 cells within the same inflammatory niche, recapitulating the cell-intrinsic licensing advantage established *in vitro* (**Fig. 4N** and **O**). Notably, this advantage was lost in both p35- and p40-deficient hosts, where T+6 and 1+6+23 cells generated equivalent and substantially reduced frequencies of IFN-γ⁺ cells with similar T-bet expression — indicating that the influence of prior polarization is muted once subsequent host cytokine signaling is disrupted. Consistent with the developmental origin of the two populations, T+6 cells retained higher IL-17A expression than 1+6+23 cells across all three host backgrounds, reflecting the persistence of TGF-β-anchored Th17 identity (**Fig. 4O**). The *p35 Rag1*^−/−^ host, which retains IL-23 but lacks IL-12, was particularly informative: although T+6 and 1+6+23 cells were equivalent, both transdifferentiated at frequencies substantially lower than in *Rag1^−/−^*hosts, yet appreciably above the *p40 Rag1*^−/−^ baseline. Thus, while IL-23 licenses Th17 cells for efficient IL-12 driven transdifferentiation, IL-12 is not absolutely required for terminal conversion: in its absence, IL-23 alone can support a partial, albeit less efficient, transdifferentiation program *in vivo* (**Fig. 4N** and **O**). The cell-intrinsic advantage conferred by early IL-23 priming therefore requires ongoing host IL-23 or IL-12 signaling to fully manifest *in vivo*: in the absence of host cytokines, the priming-derived advantage is abolished, and the two populations exhibit equivalent transdifferentiation.

Thus, our findings indicate that while IL-23 exposure during early polarization establishes a cell-intrinsic licensing state, this state is not durable without continuous reinforcement by host-derived IL-23 to enable IL-12–mediated induction of the pathogenic Th1 effector program. T+6 primed Th17 cells, having received no IL-23 during polarization, can acquire pathogenicity indistinguishable from their IL-23–primed counterparts during late development when host cytokines are available; conversely, 1+6+23 cells lose their licensing advantage when host IL-23 and IL-12 are absent. The pathogenic potential of a Th17 cell is therefore determined by cumulative IL-23 and IL-12 exposure across both its developmental history and its local tissue microenvironment, supporting a model in which IL-23 functions as a continuous and tunable licensing input rather than a one-time developmental switch — one that, even when continuously available, requires IL-12 signaling to drive full pathogenic conversion.

## Discussion

Our findings redefine the role of IL-23 in Th17 plasticity. Historically conceptualized as a terminal effector cytokine that sustains the Th17 lineage through STAT3-driven survival and proliferation, we show that IL-23 additionally has an instructive role functioning as a developmental licensor: it signals through STAT4 to upregulate IL-12Rβ2, rendering Th17 cells more competent for IL-12–driven transdifferentiation (6, 7, 24, 25). This licensing is not restricted to a single developmental window. IL-23 upregulates IL-12Rβ2 when present during early naïve T cell polarization and when introduced later to committed Th17 cells, indicating that licensing operates as a continuous and tunable process rather than a fixed developmental checkpoint. Together with the parallel upregulation of *Il23r*, which reinforces the Th17 lineage anchor, IL-23 simultaneously stabilizes Th17 identity and primes for terminal conversion — a dual function that aligns with the engagement of both STAT3 and STAT4 by the IL-23 receptor complex (4).

This temporal flexibility fundamentally challenges the dichotomy used to classify Th17 cells. Polarization with TGF-β and IL-6 has classically defined “non-pathogenic” Th17 cells, while inclusion of IL-1β, IL-6, and IL-23 generates a “pathogenic” phenotype (13, 22, 23). Our findings argue that there is nothing inherently or permanently restrictive about the non-pathogenic state. Non-pathogenic Th17 cells transferred into *Rag1*^−/−^ hosts acquire pathogenicity functionally indistinguishable from their IL-23 primed counterparts, and this acquired pathogenicity is lost in hosts lacking IL-23 or IL-12. Phenotypically “non-pathogenic” Th17 cells are therefore functionally interchangeable with their pathogenic counterparts when the host environment supplies the requisite licensing signals, and cumulative IL-23 exposure — encompassing both prior polarization history and ongoing host-derived signals — determines pathogenic potential.

Recognizing IL-23 as a continuous licensing agent invites reconsideration of classical *in vivo* models that have shaped current understanding of IL-12 and IL-23 biology. Much of the prevailing framework — that IL-23 drives local intestinal pathology while IL-12 mediates systemic inflammation — was established using the innate anti-CD40 colitis model in *Rag1*^−/−^ mice (17, 26). While this model demonstrated the inflammatory capacity of IL-23 in the colon, it interrogates an innate cytokine response that is mechanistically distinct from adoptive Th17 plasticity. Similarly, the CD4⁺CD45RB^hi^ naïve transfer model revealed that host IL-23, but not IL-12, is essential for colitis; this likely reflects the role of IL-23 in supporting primary T cell expansion in the absence of regulatory T cells, a process upstream of and distinct from the terminal transdifferentiation of committed Th17 cells (27–29). The genetic dependencies of these models further support this interpretation: RORγt is essential in naive transfer colitis whereas T-bet is required for transdifferentiation and disease induction in the Th17 transfer model (2, 30, 31). Rather than contradicting earlier findings, these contrasting dependencies illustrate that each model illuminates a distinct phase of the immune response. Our findings are consistent with the central conclusion of these studies — that IL-23 is the dominant and indispensable cytokine for Th17-driven colitis — but refine the role of IL-12 from dispensable to a significant, non-redundant amplifier of pathology. Host IL-12 deficiency markedly attenuated disease and transdifferentiation, whereas combined loss of IL-23 and IL-12 produced the most severe attenuation. IL-23 and IL-12 thus contribute hierarchically rather than redundantly — IL-23 as the indispensable licensing signal, sufficient on its own to drive partial transdifferentiation, and IL-12 as the principal trigger for effective terminal programming — a distinction that reconciles the apparent dispensability of IL-12 in earlier transfer models with the clear mechanistic role defined herein.

Our current findings also refine prior work from our own group on the molecular requirements for Th17 transdifferentiation. We previously showed that STAT4 and T-bet expression are essential for the Th17-to-Th1 transition, with *Stat4* deficiency contributing significantly to reduced colitis severity (2). Our current data extend these findings on multiple fronts. *Stat4* conditional knockout cells fail to transdifferentiate, supporting the central STAT4 dependence reported previously. Additionally, loss of either STAT4 or IL-12Rβ2 substantially reduces T-bet expression, and host deficiency of IL-12 or IL-23 similarly attenuates T-bet upregulation in transferred Th17 cells. These observations are consistent with T-bet being downstream of the IL-23–STAT4–IL-12Rβ2 axis and align with prior reports of IL-23-dependent T-bet induction in transitioning Th17 cells (3). We further refine the earlier conclusion that IL-12Rβ2 is dispensable for colitis: by co-transferring *Il12rb2*^−/−^ and WT Th17 cells, our competitive design controls for host-derived inflammation and reveals a clearer cell-intrinsic IL-12Rβ2 requirement than individual transfers could detect. Furthermore, earlier *in vitro* studies in which prolonged maintenance of Th17 cells in IL-23 generated IFN-γ–producing progeny used lower IL-23 concentrations (∼1 ng/mL) that produced incomplete lineage anchoring; together with the cumulative effects of prolonged TCR stimulation, this likely drove the gradual phenotype shift historically attributed to IL-23 itself (8). Our data demonstrate that IL-23 anchors the Th17 lineage in a dose-dependent manner while licensing cells for efficient terminal transdifferentiation upon subsequent IL-12 encounter. Importantly, transcriptional profiling from those foundational studies is consistent with our current licensing model when reexamined: *Ifng* and *Tbx21* upregulation, along with *Il17a* and *Rorc* extinguishment, tracked with IL-12 exposure rather than IL-23 itself (8).

The two-step licensing architecture identified here — wherein the licensing signal (IL-23) is divorced from the terminal trigger (IL-12) — suggests a stringent safeguard against runaway pathogenic transdifferentiation. The distinct cellular sources of these cytokines lend biological plausibility to this model: IL-23 is produced constitutively by CX3CR1⁺ mononuclear phagocytes and CD103⁺CD11b⁺ dendritic cells in the intestinal lamina propria in response to homeostatic microbial sensing, whereas IL-12 production by activated dendritic cells and inflammatory monocytes is induced mainly during acute inflammatory responses (11, 32, 33). This temporal segregation — homeostatic, microbiota-driven IL-23 preceding inflammation-induced IL-12 — creates the sequential exposure pattern predicted by our model, ensuring that Th17 cells undergo licensing in the homeostatic mucosal environment before encountering IL-12 in inflamed tissue. Consistent with this temporal logic, our finding that simultaneous high-dose IL-23 antagonizes IL-12–driven transdifferentiation indicates that the relative kinetics and stoichiometry of these cytokines may function as a quantitative rheostat — with appropriately timed exposure permitting licensing and concurrent exposure favoring Th17 maintenance. This rheostat is further tuned by TGF-β which acts in opposition to IL-23. TGF-β is a known suppressor of IL-12Rβ2, and we further find that TGF-β containing conditions selectively upregulate *Il6st* (gp130) expression in Th17 cells, enhancing IL-6 signaling capacity, and thereby biasing the cell towards sustained STAT3-driven Th17 lineage maintenance while restricting IL-12 responsiveness (34, 35). IL-23, conversely, augments IL-12 responsiveness through *Il12rb2* upregulation without compromising the STAT3 driven Th17 lineage anchor sustained via *Il23r*. Thus, the balance between TGF-β and IL-23 calibrates Th17 responsiveness to IL-12 and dictates Th17 plasticity in the tissue microenvironment.

This two-step licensing architecture likely reflects an evolutionary requirement for coordinated host defense at mucosal surfaces. Th17 cells are central to protection against extracellular bacteria, and their stepwise conversion into IFN-γ producers equips the host with an additional layer of antimicrobial defense that is engaged upon sequential IL-23 and IL-12 exposure (1). We have recently shown that intestinal epithelial cells (IECs) directly present cytosolically delivered *Citrobacter rodentium* antigens to CD4⁺ T cells, driving local clonal expansion and the formation of tissue-resident memory populations that protect the barrier upon reinfection (36). Because IFN-γ potently upregulates MHC class II on IECs and simultaneously polarizes myeloid populations toward pro-inflammatory, microbicidal effector states, licensing Th17 cells for IL-12– driven IFN-γ production would enhance both epithelial antigen presentation and innate effector function at the site of infection (37–39). In this view, homeostatic Th17 cells constitute a flexible reservoir that can be rapidly redeployed as IFN-γ producers to orchestrate epithelial and myeloid defense against invasive pathogens, with the temporal segregation of IL-23 and IL-12 ensuring that this layered program is coupled to pathogen-driven inflammation rather than constitutive microbiota sensing.

This licensing framework may extend beyond colonic inflammation. Previous fate mapping studies in EAE demonstrated that IL-17A+ Th17 cells transitioning toward IFN-γ-producing “ex-Th17” effectors selectively upregulate *Il12rb2* and *Tbx21* while downregulating *Il23r* and *Rorc*, and that this transition required IL-23 signaling (3). Notably, *Il12rb1* expression remained unchanged between the IL-17A-maintaining (CCR6+) and transitioning (CCR6−) populations (3). More recent single-cell fate-mapping studies have identified a stem-like, homeostatic Th17 reservoir (SLAMF6⁺) in the intestine that gives rise to encephalitogenic, IFN-γ–producing effectors (CXCR6⁺) during experimental autoimmune encephalomyelitis in an IL-23R– dependent manner (10). Chromatin accessibility profiling of these two populations showed that *Il12rb2* — together with downstream effectors of the IL-12/Th1 program including *Tbx21*, *Ifng*, *Ifngr1*, *Bhlhe40*, and *Csf2* — is selectively accessible in CXCR6⁺ pathogenic Th17 cells (10). The IL-12Rβ2 upregulated by IL-23 signaling in our system thus corresponds to a locus made accessible during transition from homeostatic to pathogenic Th17 cells in vivo. Although not tested directly here, our findings together with these observations raise the possibility that microbiome-driven IL-23 in the intestinal lamina propria licenses homeostatic Th17 cells by signaling through STAT4 to upregulate *Il12rb2*, enabling subsequent transdifferentiation upon IL-12 encounter in the inflamed tissue.

These observations also place IL-12Rβ2 within the broader architecture of pathogenic Th17 differentiation. The transition from a homeostatic to a pathogenic effector state involves coordinated remodeling of numerous loci, and is shaped by transcription factors including Bhlhe40, c-Maf, Blimp-1, and Bach2 (40–43). IL-12Rβ2 is one component of this broader transcriptional and chromatin landscape rather than its sole determinant. IL-12Rβ2 nonetheless occupies a mechanistically privileged position: As the receptor subunit that gates responsiveness to IL-12, it determines which extracellular signal the cell can receive and therefore which downstream transcriptional programs are engaged. IL-23-driven IL-12Rβ2 upregulation thus functions not as the entirety of the pathogenic program but as a critical upstream checkpoint that governs its execution.

Importantly, the licensing axis is dominant but not absolute. Previous studies in EAE showed that IL-12 is not an absolute prerequisite for clinical CNS autoimmunity. In its absence, Th17 cells can drive an atypical, neutrophil-predominant form of EAE mediated by IL-17 and GM-CSF (44–46). We similarly observe some residual IFN-γ production in p35- and p40-deficient hosts along with significantly attenuated but not fully abolished disease in p35-deficient recipients, indicating that IL-12 is not the universal gatekeeper of pathology and that additional pathways can sustain Th1-like differentiation independently of the IL-12/IL-23 axis. Autocrine IFN-γ amplification, which drives T-bet expression through STAT1-mediated positive feedback, and type 1 interferon signaling represent plausible candidate mechanisms (47–49). A contribution of IL-23 itself, acting in concert with these or other signals, also cannot be excluded, consistent with the partial transdifferentiation observed in p35-deficient hosts. Whether these pathways operate in parallel to or downstream of IL-23 licensing, and whether they sustain residual disease when the IL-12/IL-23 axis is therapeutically attenuated, are open questions with direct therapeutic relevance.

Several limitations warrant consideration. While the SMARTA TCR-transgenic system provides a precise antigen-specific framework that eliminates confounders of polyclonal TCR engagement, these cells recognize a viral epitope (LCMV gp61) rather than a microbiota-derived antigen (50). Whether the licensing kinetics observed here apply equivalently to lower-affinity, microbiota-reactive TCRs that drive intestinal inflammation in IBD will require further investigation. Our transfer colitis experiments, which operate independently of the SMARTA system, provide complementary validation. Our pathogenic polarization condition additionally includes IL-1β, and while our IL-23–isolating experiments establish that IL-23 signaling is sufficient to upregulate IL-12Rβ2, a contribution of IL-1β cannot be formally excluded. Furthermore, while our adoptive transfer colitis experiments clearly define the role of the IL-23 licensing axis in driving transdifferentiation of committed Th17 effectors, whether IL-23 plays a similar licensing role in developing naïve T cells or in other CD4⁺ T cell lineages remains unclear. Recent work has shown that IL-23R signaling can also endow Th1-like cells with a colitogenic transcriptional program distinct from that of Th17 cells, underscoring that IL-23R expression and function are not restricted to the Th17 lineage (51). Accordingly, our findings should not be extrapolated beyond Th17 plasticity without direct investigation in other CD4⁺ T cell subsets. Finally, direct validation of the IL-23–STAT4–IL-12Rβ2 axis in human Th17 cells will be essential to extend these findings to clinical contexts.

In summary, our findings establish a stepwise model in which IL-23 functions as a developmental licensor, signaling through STAT4 to upregulate IL-12Rβ2 and rendering Th17 cells competent for IL-12–driven transdifferentiation. By defining the IL-23–STAT4–IL-12Rβ2 axis as the molecular bridge between Th17 maintenance and pathogenic conversion, this work provides a mechanistic rationale for the clinical efficacy of IL-23–targeted therapies and identifies the licensing step as a discrete and pharmacologically tractable point of intervention (52, 53). The structure-based IL-23 mutein employed here — selectively uncoupling STAT3-dependent Th17 maintenance from STAT4-dependent pathogenic licensing — may itself represent a conceptual template for next-generation therapeutics that disarm pathogenic plasticity while preserving foundational mucosal immunity.

## Materials and Methods

### Mice

*Il17f^Thy1.1^* reporter mice and SMARTA TCR-transgenic mice have been described previously (2, 50). SMARTA-1 CD45.1, *Il12rb2*^−/−^, *Il12a*^−/−^, *Il12b*^−/−^, *Rag1*^−/−^, and *Cd4^Cre^* were purchased from The Jackson Laboratory. *Stat4*^fl/fl^ mice were a gift from Laurie Harrington (University of Alabama at Birmingham). *Il17f^Thy1.1^*reporter mice and individual knockout or transgenic strains were crossed in our facility to produce mice homozygous for the *Il17f^Thy1.1^*reporter and the respective gene deletion or transgenic alleles (e.g., *Il17f^Thy1.1^*·*Il12rb2*^−/−^; *Il17f^Thy1.1^*·SMARTA, *Stat4^fl/fl^*·*Cd4^Cre^*). All mice were on a C57BL/6 background, were *Helicobacter* spp.-negative, and were maintained under specific pathogen-free conditions. All mouse strains were bred and maintained at University of Alabama at Birmingham in accordance with the IACUC regulations.

### CD4⁺ T cell preparation and *in vitro* polarization

Naïve CD4⁺ T cells were purified from spleens of SMARTA· *Il17f^Thy1.1^* mice using the Mouse Naïve T cell isolation kit (Miltenyi Biotec) per the manufacturer’s instructions. For primary Th17 polarizations, sorted naïve CD4⁺ T cells were cultured at a 1:10 ratio with irradiated splenocyte feeders in Isocove’s modified Dulbecco’s medium containing 10% FBS, 100 IU/mL penicillin, 100 μg/mL streptomycin, 1 μM sodium pyruvate, nonessential amino acids, 2.5 μM β-mercaptoethanol, and 2 μM l-glutamine (I10 medium). SMARTA CD4⁺ T cells were stimulated with 1.25 μg/mL gp61 peptide (KAVYNFATC), anti-CD28 (1 μg/mL) and either classical Th17-inducing cytokines (2.5 ng/mL rhTGF-β1 and 10 ng/mL rmIL-6; “T+6”) or pathogenic Th17-inducing cytokines (20 ng/mL rmIL-1β, 10 ng/mL rmIL-6, and 10 ng/mL rmIL-23; “1+6+23”). For adoptive transfer experiments, similar conditions were used except that cells were stimulated with anti-CD3 (2.5 μg/mL). Cultures additionally contained 10 μg/mL anti-IFN-γ (XMG1.2) and 10 μg/mL anti–IL-4 (11B11). After 5 days, viable IL-17F^Thy1.1+^ cells were sort-purified on BD FACSAria I or II. A complete list of reagents is provided in SI Table S1.

### Secondary and tertiary cultures

IL-17F^Thy1.1+^ sorted cells were restimulated at a 1:10 ratio with irradiated *Il12b*^−/−^ splenocyte feeders in I10 medium containing 1.25 μg/mL gp61 peptide and indicated cytokine conditions: T+6 (TGF-β1 + IL-6), wild-type IL-23 (1 or 10 ng/mL), IL-23 mutein (10 ng/mL), or titrated IL-12 (0.0001–10 ng/mL). For tertiary cultures examining late-stage licensing, T+6–primed cells were first maintained for 5 days in secondary culture with T+6, WT IL-23, or IL-23 mutein. Live CD4+ cells were then sorted and re-stimulated with titrated rmIL-12 in tertiary culture for an additional 5 days, using the same TCR and co-stimulation conditions as primary and secondary cultures.

### IL-23 mutein generation and validation

The IL-23 mutein with substitutions P39A, D40A, E81A, and F82A in the p40 subunit was generated as previously described and obtained from the laboratory of K.C. Garcia (20). Selective abolition of STAT4 phosphorylation with preservation of STAT3 activation was confirmed by phospho-flow cytometry on Th17 cells stimulated with WT or mutant IL-23.

### Adoptive Th17 transfer colitis model

For individual transfers, 4 x 10^5^ IL-17F^Thy1.1+^-sorted Th17 cells (T+6– or 1+6+23–polarized) were injected intraperitoneally into age- and sex-matched *Rag1*^−/−^, *Il12a*^−/−^·*Rag1*^−/−^, or *Il12b*^−/−^·*Rag1*^−/−^ recipients. For competitive co-transfers, 2 x 10^5^ T+6 (CD45.1) and 2 x 10^5^ 1+6+23 (CD45.1/2) Th17 cells, or 2 x 10^5^ wild-type (CD45.1) and 2 x 10^5^ *Il12rb2*^−/−^ (CD45.1/2) T+6–primed Th17 cells, were co-transferred in a 1:1 ratio. Mice were monitored weekly for weight loss and signs of colitis. At 4-5 weeks post-transfer, mice were euthanized, and colons were harvested for histopathological scoring and colonic lamina propria lymphocyte isolation as described previously (2).

### Flow cytometry

For intracellular cytokine staining, cells were stimulated with 50 ng/mL PMA (Sigma) and 750 ng/mL ionomycin (EMD Millipore) for 4 hours in the presence of GolgiPlug (BD Biosciences). Cells were stained with Fixable Viability Dye eFluor 780 (Invitrogen) and surface antibodies, fixed and permeabilized with the Foxp3/Transcription Factor Staining Buffer Set (eBioscience), and stained intracellularly as described previously (2). For phospho-flow cytometry, cells were stimulated with the indicated IL-12 concentrations for 45 min at 37 °C, fixed with 1.5% formaldehyde for 15 min at room temperature, permeabilized with ice-cold 90% methanol for 30 min, and stained with phospho-STAT4 (pY693; BD Biosciences) and phospho-STAT3 (pY705; BioLegend) antibodies for 60 min at room temperature. For IL-12Rβ2 surface staining, cells were stained with anti–IL-12Rβ2-PE (BioLegend) in PBS with 2% FBS and 0.1% sodium azide for 60 min at room temperature. All flow cytometry data were acquired on Attune NxT (Thermo Fisher Scientific) and analyzed with FlowJo software (BD Biosciences). A complete list of antibodies is provided in SI Table S2.

### Quantitative RT-PCR

Total RNA was extracted using the RNeasy Mini Kit (Qiagen) and cDNA synthesis was performed using the iScript Reverse Transcription supermix (Bio-Rad) according to manufacturer’s protocol. cDNA amplification was analyzed with SsoAdvanced Universal SYBR Green Supermix (Bio-Rad) in a Bio-Rad CFX qPCR instrument. Reactions were run in triplicate and normalized to *Gapdh* and analyzed using the ΔΔC_T method. Primer sequences are provided in SI Table S3.

### Histopathological scoring

Colons were cut longitudinally; tissue from the proximal, middle, and distal portions was fixed in 10% formalin and processed for histopathological analysis. H&E-stained sections were scored by a pathologist in a blinded fashion as previously described (2, 8).

### Statistical analysis

Statistical significance was calculated using Prism software (GraphPad). The nonparametric Mann–Whitney test or Kruskal–Wallis test was used for histological scoring; all other data were analyzed using unpaired two-tailed Student’s *t* test or one-or two-way ANOVA with appropriate post-hoc tests. P values ≤ 0.05 were considered significant. See figure legends for details.

## Acknowledgements

The authors thank Laurie Harrington for providing the *Stat4*^fl/fl^ mice; members of the Weaver laboratory for helpful discussions; S. Harbour, D. Figge, and D. Pham for their intellectual contributions during this work; H. Turner, K. Cantrell, C. Shah, and S. Shah for assistance with animal husbandry; and V. S. Hanumanthu, H. C. Pal, H. Johnson, and K. Stortz at the UAB Flow Cytometry and Single Cell Core Facility. This work was supported by NIH/NCI R00CA277554 (A.S.C.), NIH/NIAID R37AI169864 (C.T.W.), and NIH T32 and F30 support (B.F.F.).

## Author Contributions

A.S.C. and C.T.W. conceived the project and wrote the manuscript. A.S.C., A.E., C.L.Z and C.T.W. performed the experiments and/or interpreted the results. B.F.F. provided technical expertise and contributed to data interpretation. Y.N.-K. conducted *in vitro* fertilization procedures to maintain all mouse lines with the same C57BL/6 microbiota. T.R.S. provided histopathologic expertise and disease scoring. C.R.G. and K.C.G. provided IL-23 muteins and helpful discussions.

---

**Figure S1.**
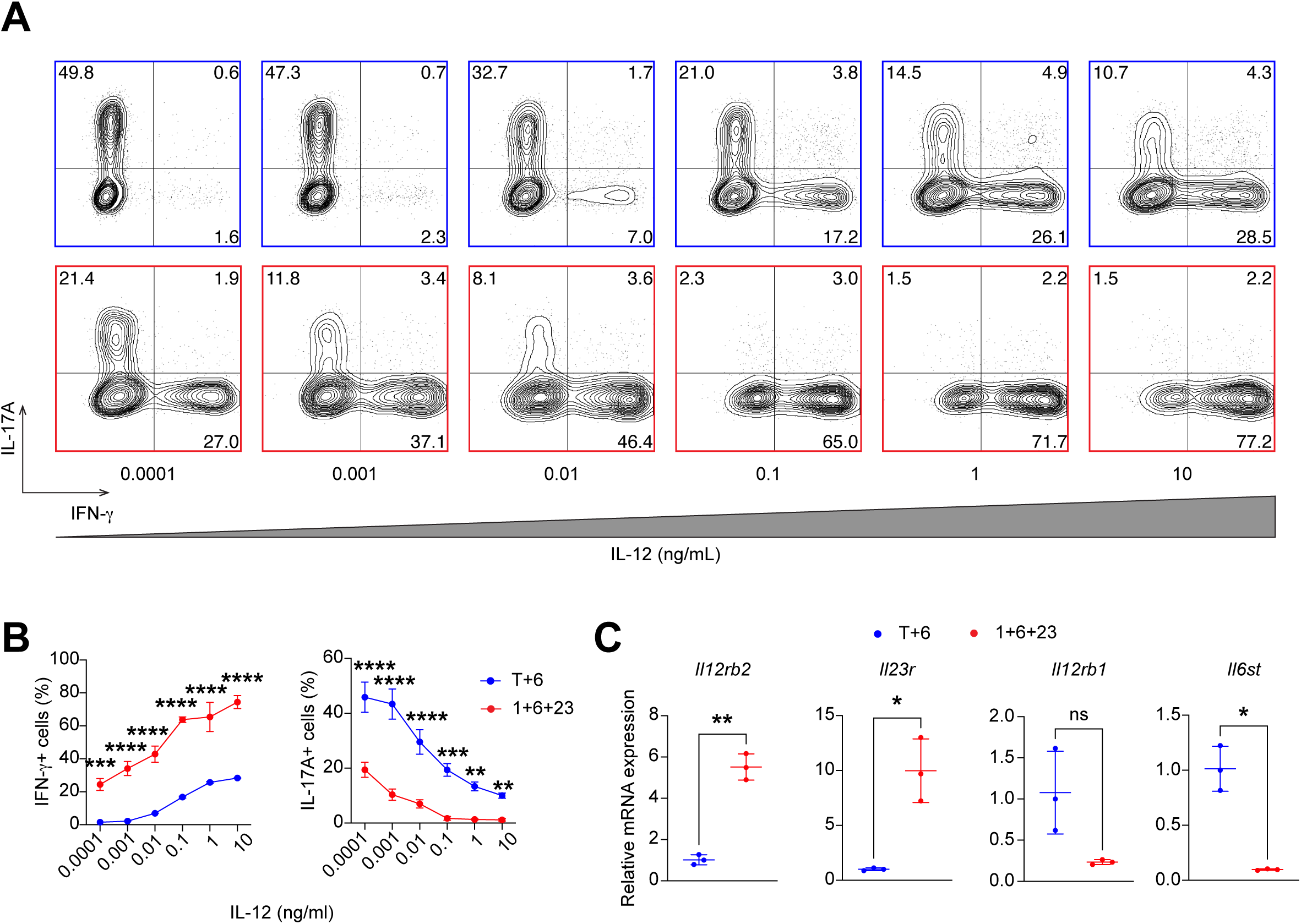
IL-23 priming lowers the IL-12 threshold for transdifferentiation and selectively upregulates cytokine-specific receptor subunits. (*A*) Representative FACS plots of IL-17A and IFN-γ expression in T+6– and 1+6+23–primed Th17 cells restimulated across a graded IL-12 dose range (0.0001–10 ng/mL) in secondary culture. Plots gated on live CD4⁺ T cells. (*B*) Frequencies of IFN-γ⁺ and IL-17A⁺ cells across IL-12 doses in T+6– and 1+6+23–primed cells (mean ± SD). (*C*) *Il12rb2*, *Il23r*, *Il12rb1* and *Il6st* transcript quantification by qRT-PCR in T+6– and 1+6+23–primed Th17 cells, normalized to *Gapdh* (mean ± SD). Data are representative of three independent experiments with three biological replicates per condition. *P < 0.05, **P < 0.01, ***P < 0.001, ****P < 0.0001; ns, not significant.

**Figure S2.**
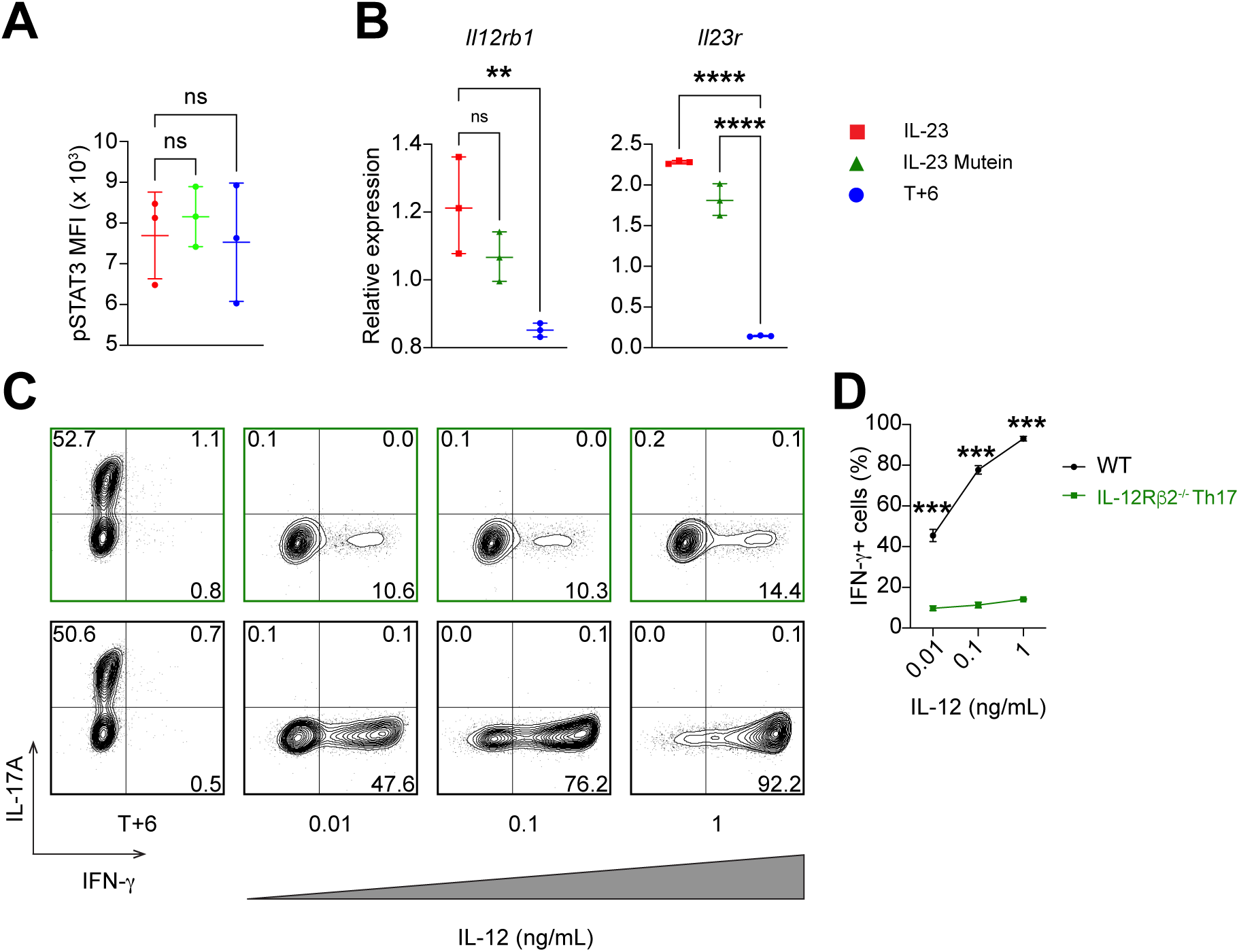
The IL-23 mutein preserves STAT3 signaling, and IL-12Rβ2 is required for IL-12–driven transdifferentiation in vitro. (*A*) pSTAT3 MFI in Th17 cells maintained in WT IL-23, IL-23 mutein, or T+6 during secondary culture (mean ± SD). (*B*) *Il12rb1* and *Il23r* transcript quantification by qRT-PCR in WT IL-23–, IL-23 mutein–, and T+6–maintained Th17 cells, normalized to *Gapdh* (mean ± SD). (*C*) Representative FACS plots of IL-17A and IFN-γ expression in *Il17f^Thy1.1^* (WT) and *Il17f^Thy1.1^*·*Il12rb2*^−/−^ Th17 cells (T+6–primed) restimulated with T+6 or titrated IL-12 (0.01, 0.1, 1 ng/mL) in secondary culture. Plots gated on live CD4⁺ T cells. (*D*) Frequencies of IFN-γ⁺ cells across IL-12 doses in WT and *Il12rb2^−/−^* Th17 cells (mean ± SD). Data are representative of three independent experiments with three biological replicates per condition. *P < 0.05, **P < 0.01, ***P < 0.001, ****P < 0.0001; ns, not significant.

**Figure S3.**
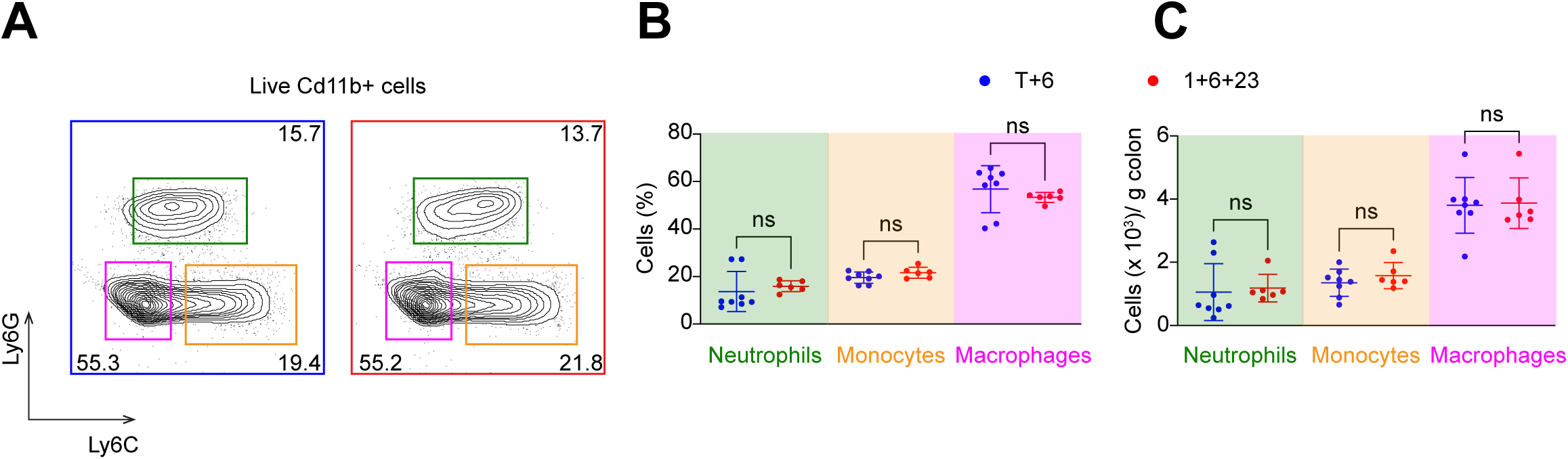
Colonic myeloid populations are equivalent in recipients of T+6– and 1+6+23– polarized Th17 cells. (*A*) Representative FACS plots of Ly6G and Ly6C expression on live CD11b⁺ myeloid cells isolated from the colonic lamina propria of *Rag1^−/−^* recipients 5 weeks after transfer of T+6– or 1+6+23–polarized Th17 cells. (*B* and *C*) Frequencies (*B*) and absolute numbers per gram of colonic tissue (*C*) of neutrophils (Ly6G^+^Ly6C^int^), monocytes (Ly6G⁻Ly6C^hi^), and macrophages (Ly6G⁻Ly6C^-^) (mean ± SD). Data are pooled from two independent experiments with 6–8 mice per group. ns, not significant.

**Table S1:** Reagents used for Th17 cultures.

| Reagent | Catalog# | Vendor |
| --- | --- | --- |
| Hamster Anti-Mouse CD3ε | Clone: 145-2C11 | BD Biosciences |
| Hamster Anti-Mouse CD28 | Clone: 37.51 | BD Biosciences |
| Human TGF-beta 1 Recombinant Protein | 240-B-010 | R&D Systems |
| *Invivo*Mab anti-mouse IFN-γ | BE0055 | Bioxcell |
| *Invivo*Mab anti-mouse IL-4 | BE0045 | Bioxcell |
| LCMV GP (61-80) | AS-64851 | Anaspec |
| Recombinant mouse IL-6 | 406ML200 | R&D Systems |
| Recombinant mouse IL-12 | 419-ML-050 | R&D Systems |
| Recombinant mouse IL-23 | 1887-ML-010 | R&D Systems |
| Recombinant mouse IL-1β | 401ML025 | R&D Systems |

**Table S2:**
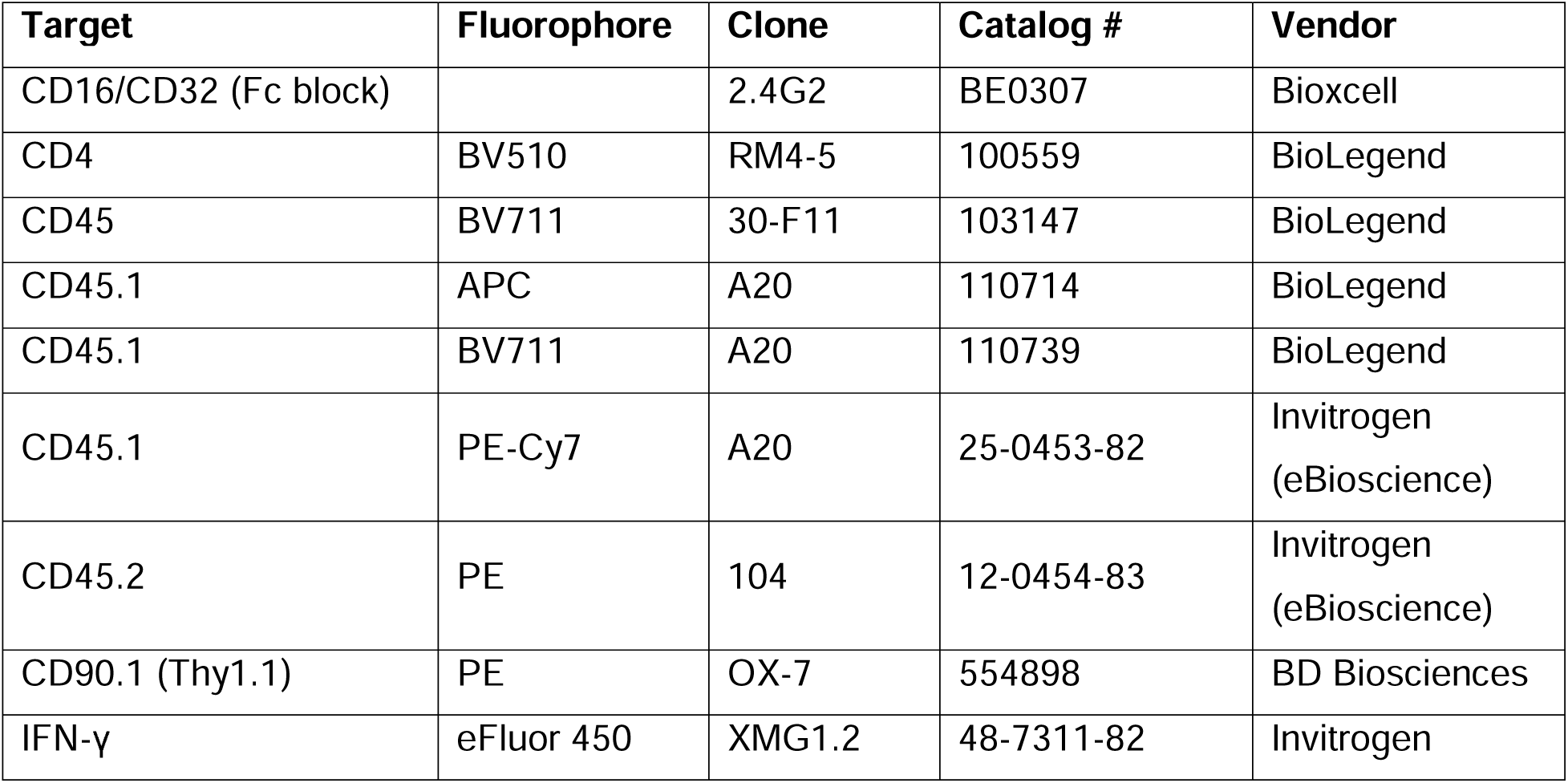
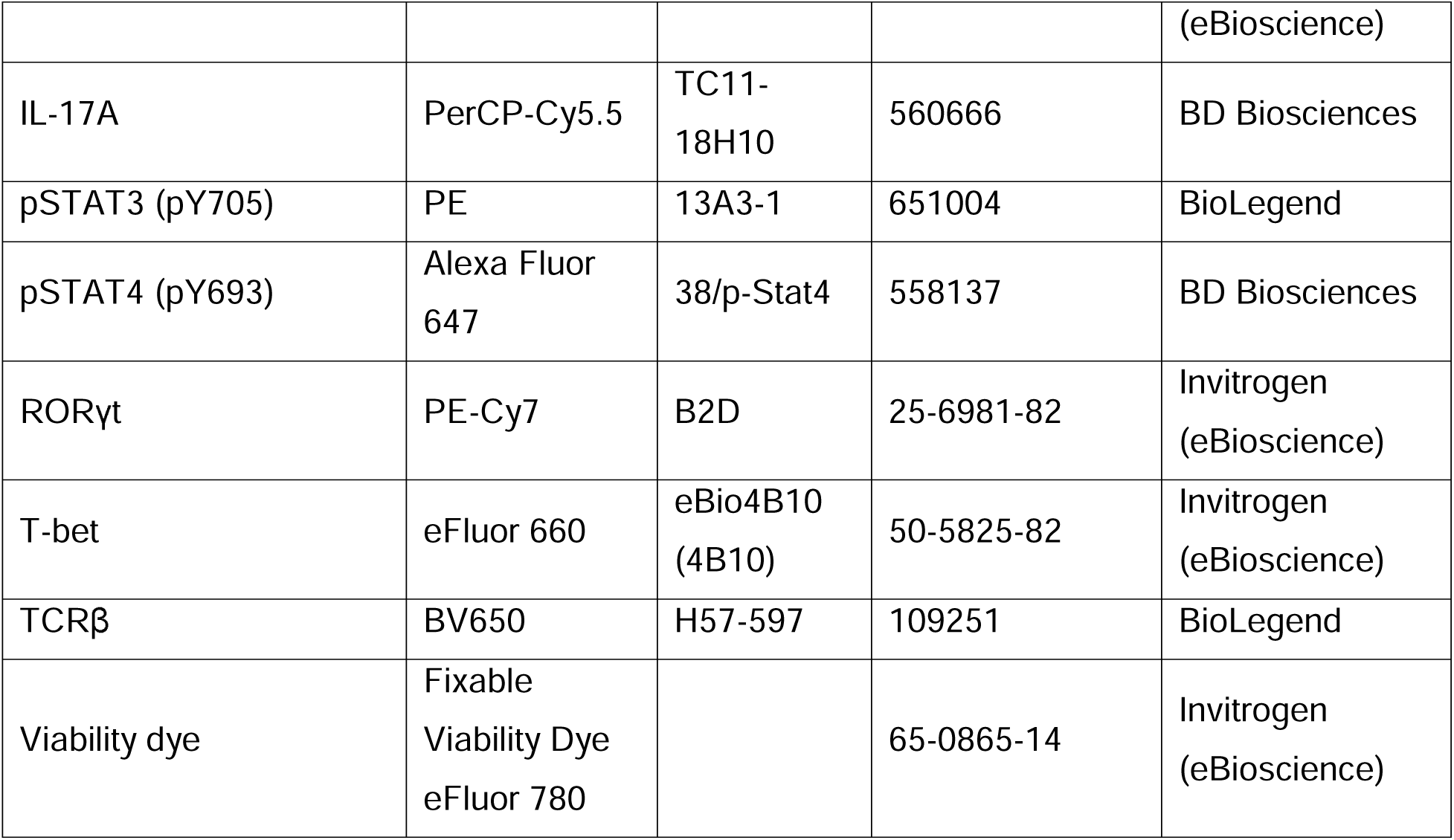
Antibodies used for surface and intracellular staining.

**Table S3:**
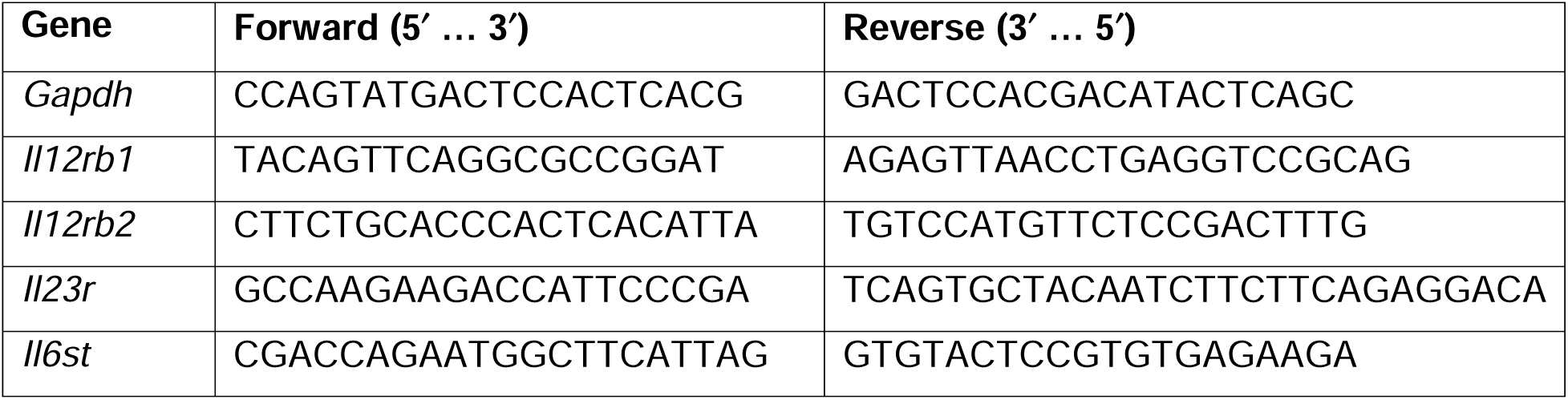
qRT-PCR primer sequences.

